# Crab fisherman communities in north Brazil: a new high risk population for vampire bat rabies

**DOI:** 10.1101/590083

**Authors:** Nailde de Paula Silva, Elane Araújo Andrade, Denis Cardoso, Ruth de Souza Guimarães, Mateus Borges Silva, Kelly Karoline Gomes Nascimento, Diego de Arruda Xavier, Isis Abel

**Affiliations:** Laboratory of Epidemiology and Geoprocessing (EpiGeo), Veterinary Medicine Institute, Federal University of Pará (UFPA), Castanhal, Pará, Brazil.; Veterinary Medicine Department, Federal University of Lavras (UFLA), Lavras, Minas Gerais, Brazil

## Abstract

An outbreak of human rabies transmitted by hematophagous bats occurred in 2018 in the state of Pará, Brazil, eastern Amazon, after 14 years with no record of the disease. It is necessary to understand the epidemiological characteristics of these attacks to protect the local population. This study aimed to characterize attacks of humans by vampire bats in the municipality of São João da Ponta, Pará state, Brazil, from 2013 to 2015. All individuals attacked by bats who sought medical care during the study period (n=5) were identified in the Notifiable Diseases Information System (SINAN) database and answered a questionnaire about the circumstances of the attack. Using snowball sampling, seed cases identified other individuals who were attacked in the same period but did not seek medical care (n=61), totalizing 66 people attacked in the same period. The interviewees were male (92.4%), adults between 20 and 50 years old (69.6%) and had completed elementary education (86.3%). Most were rural residents (92.4%) and crab fishermen (79.3%). The interviewees (92.4%) identified the mangrove of the Mãe Grande de Curuçá extractive reserve as an area conducive to attacks by vampire bats, where groups of fishermen sometimes concentrate for days for crab fishing, often living in improvised dwellings without walls and covered by tarps or straw (88.8%). The wounds were single bites (71.2%) and were located on the lower limbs (93.9%). Overall, 42.4% of participants had been bitten more than four times throughout their life (range 1-23 attacks). Participants were unaware of the risk of contracting rabies by the bite (95.4%). Using São João da Ponta as a model, this study shows that bat attacks are an essentially occupational problem in the study region. Indeed, for each reported attack, there are 12.2 unreported cases. It is necessary to develop strategies to reach this population for prophylactic treatment.

**Author Summary:** Different from which occurs worldwide in relation to rabies transmission, in Amazon region, vampire bat is involved on direct transmission of rabies virus to humans when searching for bloodmeal. It is common in the state of Pará, Eastern Amazon, large areas inhabited near forests and mangroves. People living there use forest natural resources as a way of income and sustenance and these working conditions is what our study points out as an important factor for aggressions predisposition. Here this subject is shown as an occupational problem. This study also quantified for the first time underreported human’s aggressions by bats in Amazon, using the snowball sampling, which valued the relationship between individuals to reach the target population. Based on these results, rabies surveillance may direct actions for prevention and health education for these individuals, including changes in notifications forms and suggesting pre-exposure prophylaxis in vaccination calendar of the Brazilian Ministry of Health for these individuals exposed to the rabies virus.

## Introduction

In general, the causal factors of rabies are manifold. The epidemiological cycle of the viral agent is complex and involves several mammalian hosts. Many researchers consider contact with the wild reservoir, human population mobility, and the social and cultural characteristics of a population as the main determinants for onset of the disease. The high impact of rabies transmission by vampire bats in the Amazon region is unquestionable [1–3].

Climate, seasonality, proximity to the forest, and the presence of livestock and natural prey provide conditions conducive to the proliferation of vampire bats in the Amazon. However, this in itself does not determine the occurrence of bites and transmission of rabies in humans. It is the relationships between humans and the environment that place human populations at epidemiological risk, which is also associated with the lack of medical care, poverty and low level education [4,5].

This relationship between humans and the environment is reflected in the use of the mangroves’ natural resources by the families in that region. The municipality of São João da Ponta, Pará state (PA), is located in an Extractive Reserve (São João da Ponta RESEX) in the Eastern Amazon and is surrounded by other conservation units. The main economic activity of the municipality is swamp ghost crab (*Ucides cordatus*) fishing [6]. Informal conversations with the population indicated that many residents in the municipality had been attacked by bats but did not seek medical care.

According to the guidelines of the Health Surveillance Department of the Ministry of Health (SVS/MS), bat bites are subject to mandatory notification throughout Brazil. Every case of a bite or attack involving bats must be notified to health authorities using the Notifiable Diseases Information System (SINAN) and completing the human anti-rabies treatment notification form [7]. This form collects identification data of the notifying unit and agent, patient identification, characteristics of the injury, characteristics of the animal responsible for the attack (in cases involving dogs or cats), place of residence and treatment regimen. This form must be completed at each visit and sent to the appropriate department for processing. Although this form has a lot of information to aid in rabies surveillance, it does not provide a record of the circumstances of the attack, which is a key for the surveillance of cases in which the attacking animal is a vampire bat.

To help in the recognition of areas where populations are susceptible to vampire bat attacks, and assuming that the population of the municipality of São João da Ponta is representative of the crab fishermen riverside communities in Amazon RESEX area study attacks by vampire bats in mangrove areas of the Eastern Amazon, Pará, Brazil.

## Methods

### Ethical considerations

This study was conducted with the authorization of the National Research Ethics Commission (Portuguese acronym: CONEP) of the National Health Council (Portuguese acronym: CNS; registration number CAAE: 49593315.1.0000.0018), the Brazilian Institute of the Environment and the Chico Mendes Institute for Biodiversity Conservation (Portuguese acronym: ICMBio; registration number 50344-1). All study participants provided written informed consent.

### Description of the study area

The Salgado micro-region in Pará comprises 11 municipalities that are home to five Marine Extractive Reserves (RESEX): São João da Ponta RESEX, where the municipality under study is located, Maracanã RESEX, Mocapajuba RESEX in São Caetano de Odivelas, Mestre Lucindo RESEX in Marapanim, and Mãe Grande de Curuçá RESEX. The latter covers practically the entire territory of the municipality of Curuçá-PA. The climate is hot and humid, with a rainy season (from January to June) and a dry season (from July to December) [8][9]. The vegetation is composed of moderately preserved mangrove forest with species such as *Rhizophora mangle, Avicennia germinans* and *Laguncularia racemosa* [10](Figure 1).

**Fig 1.**
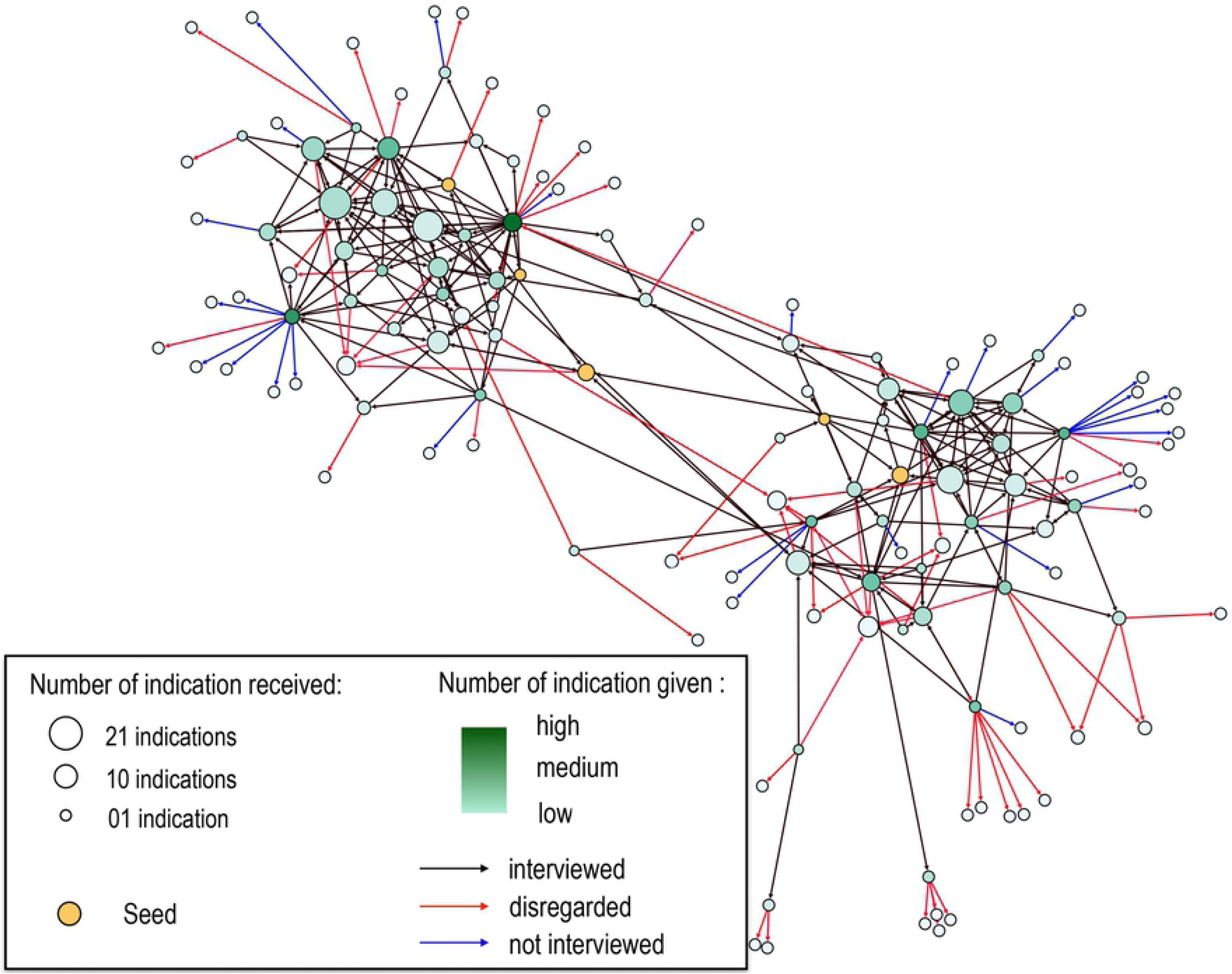
Study area. Distribution of extractive reserves in the Salgado microregion, Pará, Brazil. The map was created specifically for this manuscript and was generated by ArcGis 10.1 (ESRI) based in Brazilian Institute of Geography and Statistic Database (https://downloads.ibge.gov.br/)

### Data collection and analysis

Information of individuals attacked by bats in São João da Ponta municipality between 2013 and 2015 were from Notifiable Diseases Information System (SINAN). This period was selected to prevent the interviewee from having difficulty remembering the episode and answering the questions.

These individuals were visited in their homes, and during these visits, the study objectives were explained, and the informed consent form was signed. Subsequently, a semi-structured questionnaire was used to collect data on the circumstances of the attack, such as location and time of the attack, type of attack, time elapsed between attack and anti-rabies treatment, frequency of attack, professional occupation of the attacked individuals and structure of the household or shelter where the attack occurred.

A non-probabilistic snowball sampling technique was used to obtain information on individuals who were attacked by bats during the same period but did not seek medical care. The seeds were the cases reported in SINAN during the study period [11]. Individuals reached by this method were also interviewed and all locations indicated by the interviewees as the area of the attack were georeferenced.

The descriptive statistical analysis of the data was performed using SPSS v. 20. A geographic database was created with the coordinates of the locations of the attacks. Using the cartographic databases of the Brazilian Institute of Geography and Statistics (IBGE), the spatial distribution of cases was analyzed in ArcGIS™ 10.1. The relationships between the individuals reached by snowball sampling was visualized using R.

## Results

Between 2013 and 2015, five residents of the São João da Ponta municipality sought anti-rabies treatment after being attacked by vampire bats and therefore were registered in SINAN. These individuals identified another 141 individuals who were attacked by bats in the same period. Of those, 61 were interviewed and the others 80 individuals were disregarded in the analysis because they no longer lived in the community or were bitten for more than three years or refused to answer the questions and we could not confirm the aggression. Therefore, for each individual who sought care, there were at list 12.2 cases that were not reported. The communication network between these individuals is illustrated in Figure 2. We can observe that there are individuals who have more indications. They are individuals better known in the community. In contrast, there are few individuals who know more individuals who have been attacked by bats. To these, besides being more connected with the others in the network, they have high “PageRank” (centrality measures that reflects how much is indicated by the others of the network and those that indicate also have greater popularity). So, we can admit that people have neighbors that have more indications than them, corroborated by the article of Feld (1991) [12]. One of these more connected individuals was a Community Health Agent, a professional that visit households daily as a municipality action of preventive medicine.

**Fig 2.**
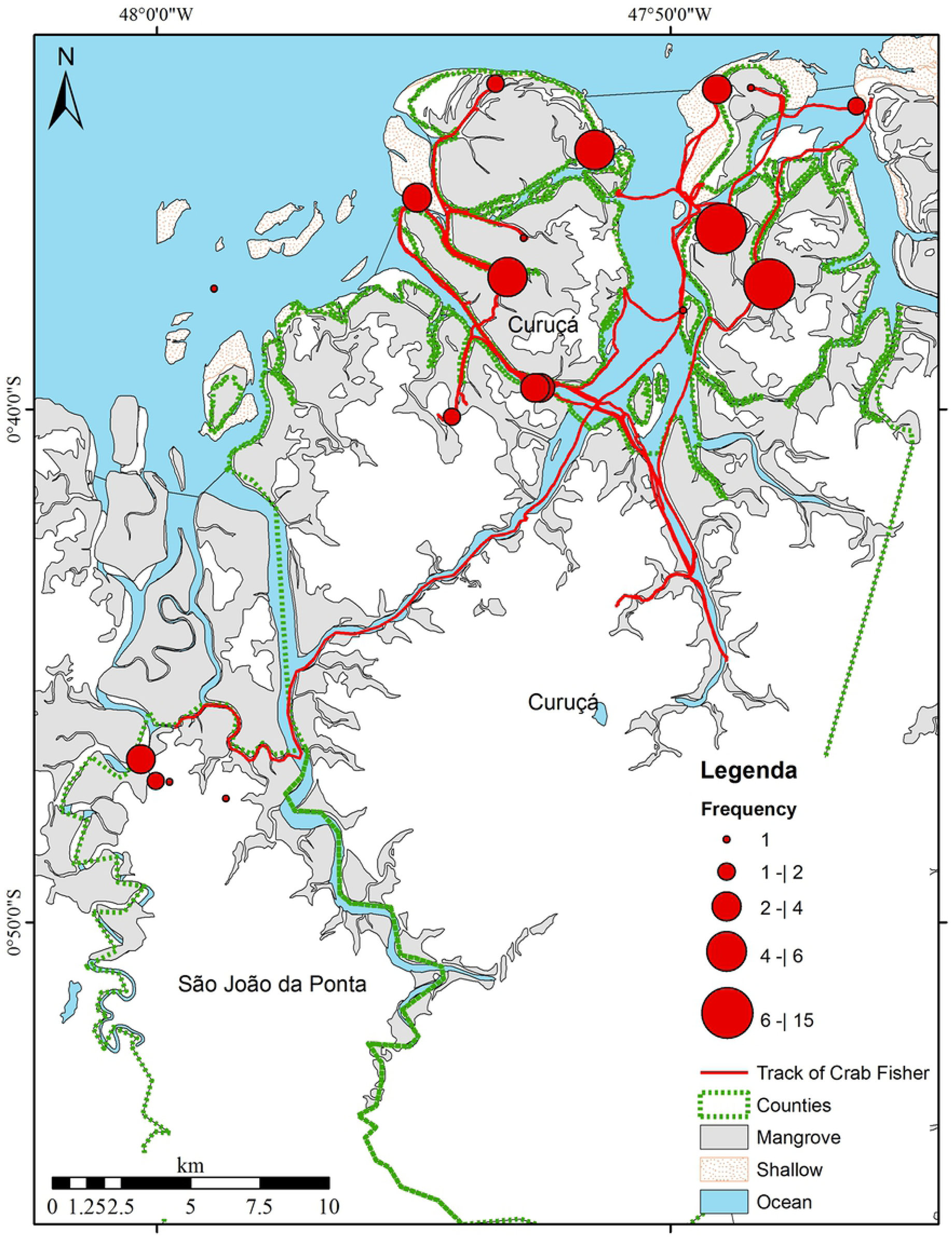
Network of relationships between the residents of the São João da Ponta municipality (Pará, Brazil) who were attacked by bats between 2013 and 2015. The nodes represent the individuals who were attacked. The diameter of the nodes is proportional to the number of times the person was mentioned.

Among the victims, 10 (15.1%) reported that the attacks occurred in their homes or in areas near the São João da Ponta Extractive Reserve, whereas the majority (84.9%) were attacked in the mangrove areas of the Mãe Grande de Curuçá Extractive Reserve in Curuçá municipality, a neighboring city (Figure 3).

**Fig 3.**
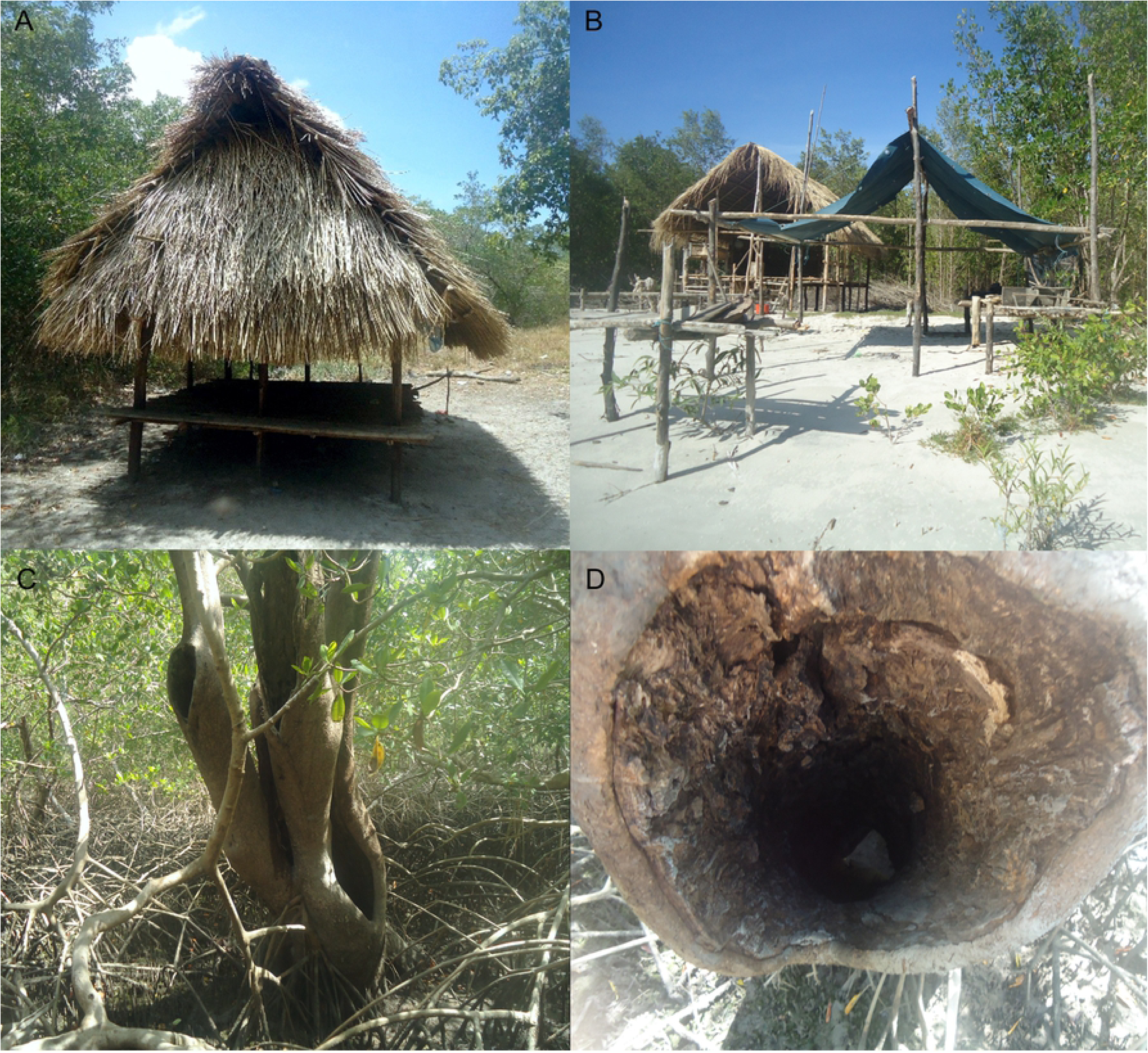
Spatial distribution of locations where residents of the Curuçá municipality were attacked by bats. The lines indicate the route traveled by the crab fishermen from their municipality of origin to the place where they camped for crab fishing. The map was created specifically for this manuscript and was generated by ArcGis 10.1 (ESRI) based in Brazilian Institute of Geography and Statistic Database (https://downloads.ibge.gov.br/)

Nineteen sites were identified as locations of attacks: 14 in the Curuçá RESEX, four in the São João da Ponta RESEX and one in the Marapanim RESEX. The Cuimiri and Pacamorema beaches which are both located in Curuçá, accounted for the highest proportions of bites, 22.7% and 10.6% respectively. Figure 4 shows the route traveled by the crab fishermen between the locations of the attack and their city of origin, where they would seek anti-rabies treatment.

**Fig 4.**
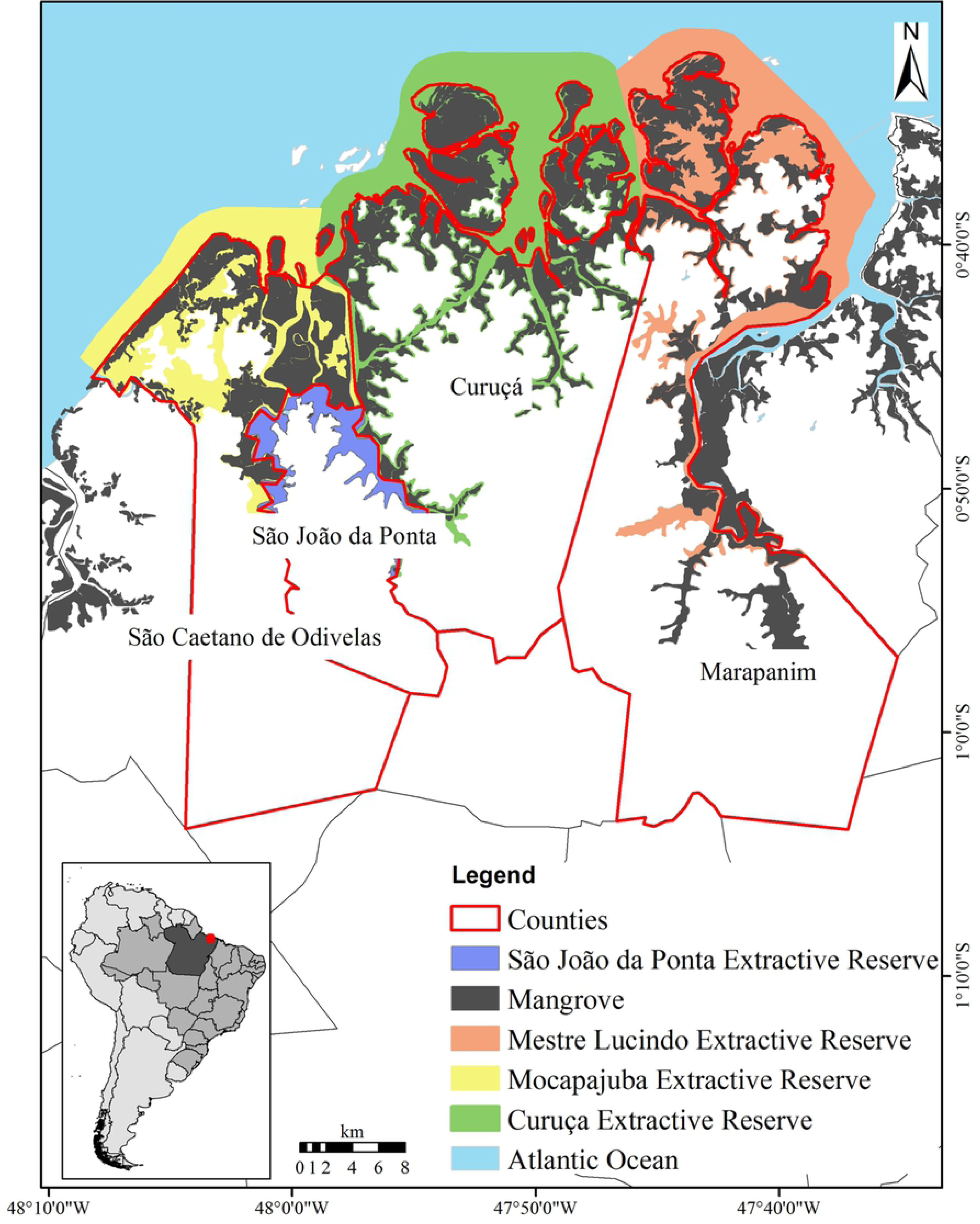
Housing conditions of the individuals who were attacked. A) shack without walls and with a straw roof; B) Shack without walls and with a tarp roof; C and D) Siriubeira.

The attacked individuals were mostly male (92.4%) and adult (69.6%) and had less than 4 years formal school education (54.4%). Most lived in the rural area of the city (92.4%) and were crab fishermen (79.3%) (Table 1).

**Table 1.** Characteristics of individuals bitten by bats in the São João da Ponta municipality, Pará, Brazil.

| VARIABLES | RESPONSE IN DESCENDING ORDER OF FREQUENCY (%) |  |  |
| --- | --- | --- | --- |
|  | 1<sup>a</sup> | 2<sup>a</sup> | 3<sup>a</sup> |
| **CHARACTERISTICS OF GRAZING** |  |  |  |
| **Ocupation** | Crabfishing (79.3) | Student (12.1) | Farmer (6.1) |
| **Sexo** | Male (92.4) | Female (7.6) | - |
| **Age** | 20 to 50 (69.9) | 10 to 20 (28.8) | less than 10 (1.5) |
| **CHARACTERISTICS OF THE PLACE OF AGGRESSION** |  |  |  |
| **Type of home** | Tent or straw taps (88.8) | Other (without wall) (10.5) | Masonry (with wall and ceiling) (7.5) |
| **Are there animals at the scene of the aggression?** | Wild (75.8) | Domestic (48.5) | - |
| **How many people were at the scene of the aggression?** | More than five (63.3) | Two to five (24.1) | One (4.5) |
| **Was there any light source in place?** | Yes (87.9) | No (12.1) | - |
| **There is fresh water on site?** | Yes (59.1) | No (40.9) | - |
| **Where were you bitten??** | Beach (56.1) | Mangrove (30.3) | “Tiso” (3.0) |
| **In what circumstances have you been bitten?** | Fishing (83.1) | At leisure (6.2) | Others (6.2) |
| **At what time of year?** | Seca (54.5) | Chuvosa (45.5) | - |
| **FEATURES OF THE INJURY** |  |  |  |
| **How many times have you been bitten in the last 3 years?** | One (33.3) | More than five (25.8) | Two (21.2) |
| **When was the last aggression?** | 1 to 3 anos (37.9) | 2 to 6 months (34.8) | 7 months to 1 any (16.7) |
| **What was the amount of bites in the last attack?** | More than tree (45.5) | One (33.3) | Two (21.2) |
| **Type of Injury** | Single (71.2) | Multiple (28.8) | - |
| **Anatomical location of the wound** | Lower member (80.8) | Head (12.8) | Upper limb (5.13) |
| **Did you look for the health unit?** | No (86.4) | Yes (13.6) | - |
| **Do you believe that the bat bite causes any disease?** | Yes (80.3) | No (19.7) | - |
| **Do you know what rabies is?** | No (94.5) | yes (5.5) | - |

The interviewees reported that during crab fishing, they gather for days in makeshift dwellings without walls and covered by tarp or straw (88.8%) (Figure 4). In most cases, these shacks were set up in drier areas within the mangrove (30.3%) or on beaches (56.1%), less than five kilometers from the forest (73.3%) and freshwater bodies (45.4%). In addition, many fishermen reported that bats sheltered in a tree typical of the mangrove area, commonly known as black mangrove (*Avicennia germinans)* (80.3%). Most of the interviewees reported the presence of domestic animals like dogs (56.5%), domestic birds (23.9%) and bovine (10.9%) where attacks occurred. Wild animals were also observed (75.8%), with raccoon (*Procyon cancrivorus)* (26.1%), scarlet ibis *(Eudocimus ruber)*, great egret (*Ardea alba)* (23.5%) and monkey (19.1%) being the most commonly reported species. Some interviewees had found lesions suggestive of bat attacks on dogs (42.9%) and domestic birds (10.6%).

On the nights that the attacks occurred, most of the individuals (63.6%) were in a group with more than five fishermen in the same shack (ranging from 2 to 12 people), and 33.3% reported that others were also attacked the same night. The shelters typically had a light source (87.9%) such as a lantern, a bonfire or an oil lamp that was usually extinguished by strong beach winds throughout the night.

When questioned about the frequency of attacks, most reported having been attacked more than four times. The number of attacks by bats during the life ranged from 1 to 18. In most cases (34.8%), the most recent episodes had occurred between 2 and 6 months prior to the interview, more frequently in the drier season (54.4%). In general, the injuries were single bites (71.2%) and were located on the lower limbs (80.8%). Among those who sought care (i.e., the seeds, 13.6%), all completed the prophylactic regimen with 5 doses of vaccine and serum.

Questions addressing respondents’ perceptions regarding the risks associated with bat bites and their consequences revealed that most individuals did not use any type of personal protection equipment against bat attacks (63.6%). Those who claimed to use some type of protection used mosquito netting, “sapato de mangue” (resistant fabric wrapped around their feet), bonfires, or fishing nets around the shack as a physical barrier. Most (80.3%) were unaware of the risk of rabies transmission through a bat bite or even unaware of rabies (94.5%). Of those attacked who did not seek medical care, 66.7% reported “not caring about what happened", 13.6% did not know how to respond, and 7.6% mentioned the distance from the healthcare unit.

## Discussion

This is the first study to quantify the underreporting of bat attacks in humans in the Amazon using the snowball sampling method, which capitalized on relationships between individuals to reach the target population. Network analysis was useful in identifying the people most commonly mentioned by the community in relation to bat attacks, and it can be applied to identify key individuals who can be the focus of more intense health surveillance activities. People identified by the network can be great allies in implementing interventions to change habits and attitudes that may hinder prevention. Indeed, it can be useful to optimize Communities Health Agents job, when bitten people active search is necessary.

Human behavior, in particular the lack of knowledge in this study population regarding the predictable consequences of vampire bat bites hinders the use of SINAN quantitative reports as a reliable data source to establish public health policies for rabies transmission in these areas. This is because the number of people attacked by bats in this region is much higher than that recorded in the system. Moreover, the reports do not include the locations of attacks or the bat species involved. Although this characterization was carried in municipality of Pará state, São João da Ponta, we believe that it is reflective of the situation in other municipalities of the Salgado microrregion, which have similar geographic and cultural characteristics.

Strategies for rabies prevention in vulnerable populations in Amazon regions have been previously reviewed in the literature [1,3,4]. In fact, it has already been proposed that the SINAN form be changed to include fields related to dog and cat behavior to support animal rabies surveillance and prophylactic measures [13]. Although these changes have not yet been incorporated, this study demonstrates the need to adapt the form to include bat attacks, given that animal attacks relevant to rabies transmission in the Salgado microregion and the whole Amazon typically involve different species than in other regions [14]. The fact that these people are generally not bitten in their homes but, rather, in other circumstances demonstrates that the inclusion in the SINAN form of a field to report the location of attacks is necessary to better prevent vampire bat bites.

This study also shows for the first time that living conditions in the mangrove are important factors in bat attacks in the Amazon. Respondents reported that in these areas, attacks by bats have been occurring for a long time and have never resulted in death. In fact, there are no reports of human deaths from the rabies virus in the study region, and there were no reported cases of neurological syndromes in livestocks between 2004 and 2013 [23]. However, there are no studies on the circulation of rabies virus in this micro-region and the possibility of inexistent or inadequate communication between healthcare and agricultural services cannot be ruled out [16]. Therefore, based on the evidence of human attacks reported here and the underreporting of bites, the study region cannot be considered as an area of controlled rabies transmission. When in doubt about whether the virus is circulating, it would be prudent to define a policy of immunization and health education for this population, given the history of rabies in Pará and the prevalent risk factors for rabies [24].

Travassos da Rosa et al. (2006) stated that the proximity of animal husbandry to the dwellings was a predisposing factor for the attacks on humans that occurred in 2004/2005 in Pará state. However, these characteristics were not observed in the present study. For the population, RESEX’s are associated to the improvement of the quality of life of people that use forest resource for subsistence, being prohibited the creation of livestock in these areas, because it contradicts its rules of sustainable use [22]. These units were created for the protection of mangrove ecosystems, which constitute an area under strong anthropic pressure, with increased exploitation of their natural resources [21]. Informal reports reveal that attacks on humans increased when bovine were removed from such areas. It is also possible that hunting activities by the native population are promoting a reduction in the supply of natural prey for bats, such as the crab-eating raccoon (*Procyon carnivores*) and nonhuman primates. These animals are targeted and slaughtered by fishermen because they feed on the crabs caught in their traps, thereby reducing their daily catch. Another relevant factor is the fact that the vegetation of the area is predominantly composed of natural fields and mangroves, where little food is available for the vampire bat [20].

The age group most affected in this study differs from that reported in previous studies. Whereas most of the victims in this study were male adults, Travassos da Rosa et al. (2006) reported children as the main victims of the rabies virus transmitted by vampire bats in the 2004-2005 period. In contrast to our study, the victims in the Travassos da Rosa study were bitten in their own residences, which commonly house families with individuals of various age groups. In this context, bats seem to prefer children. In the present study, the targets of attacks were clusters of adults who gathered for work, often associated with alcohol consumption. Individuals become passive and docile victims after falling asleep following intense work activity during the day. This is similar to the behavior of herd animals, which are favored targets of vampire bats, as they seek abundant prey and tend to remain in the same place for prolonged periods to feed [15,16].

This study also demonstrates for the first time the occupational nature of the bat attack problem in the region. The vulnerability of crab fishermen is closely tied to their work activity, as they are bitten primarily while working in the mangroves. Usually these individuals dwell in shacks (huts) built on upland soil inside the mangrove or on the beaches. These dwellings are completely open, with just a tarp or straw roof, and are used for sleeping during the work period. Even students who were bitten reported that they were accompanying their parents to help with crab fishing during the school vacation.

Residents of the Deolândia, Guarajuba and Porto Grande communities accounted for most respondents. These communities are the main centers for swamp ghost crab fishing in the region, and ghost crabs are the main natural resource exploited as a source of income by these communities. Therefore, contact between humans and bats is inevitable, as crabs are the only source of income for many of these fishermen. However, the lack of individual protection equipment certainly contributes to the increased number of bat attacks in this population [2,4].

## Conclusions

Using a single city as a model for investigating bat attacks in the Amazon, this study showed that, for each person who seeks anti-rabies treatment after being attacked by bats in the region, there are 12.2 people who do not seek care and can be easily reached by surveillance agencies through snowball sampling. We also found that the population most at risk is that of crab fishermen, suggesting that bat attacks are an occupational hazard. These results can help health surveillance agencies to establish measures for the prevention of human rabies in these individuals. They can also help minimize costs and increase the efficiency of public health measures, thus avoiding the emergence of rabies in the Salgado micro-region of Pará state.

## Acknowledgments

We thank PAPQ/PROPESP-UFPA for financial support; the Secretaria Municipal Saúde de São João da Ponta for sharing the database; Conselho Nacional de Desenvolvimento Cientifico e Tecnologico (CNPq) for the research grants and the researchers JG Barreto and CCG Moraes for their valuable contributions.

